## Supplementary Information for "High-Fidelity Neural Speech Reconstruction through an Efficient Acoustic-Linguistic Dual-Pathway Framework"

Li *et al*

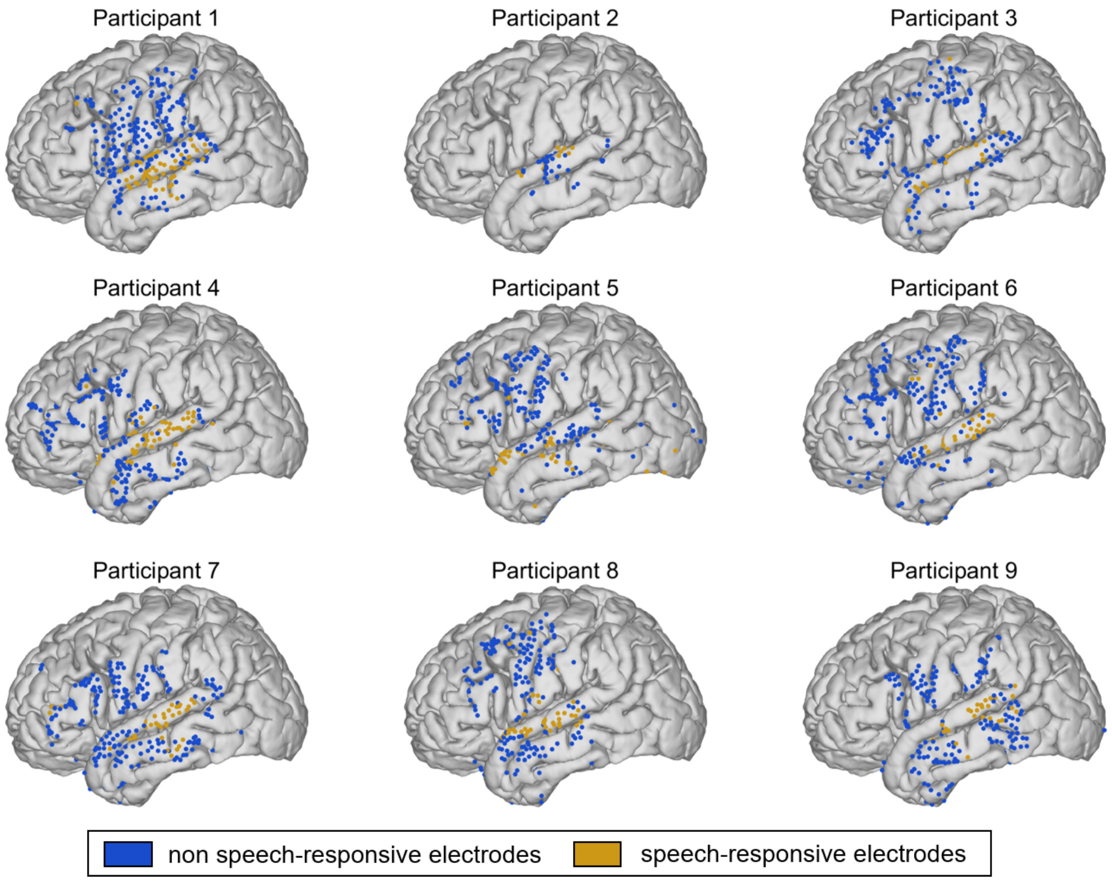


**Supplementary Figure 1. Speech responsive and tone discriminating electrodes for all participants.** ECoG grids covering the lateral temporal lobe of all participants were warped onto the MNI152 template. Yellow electrodes are responsive to speech, while blue electrodes are not.

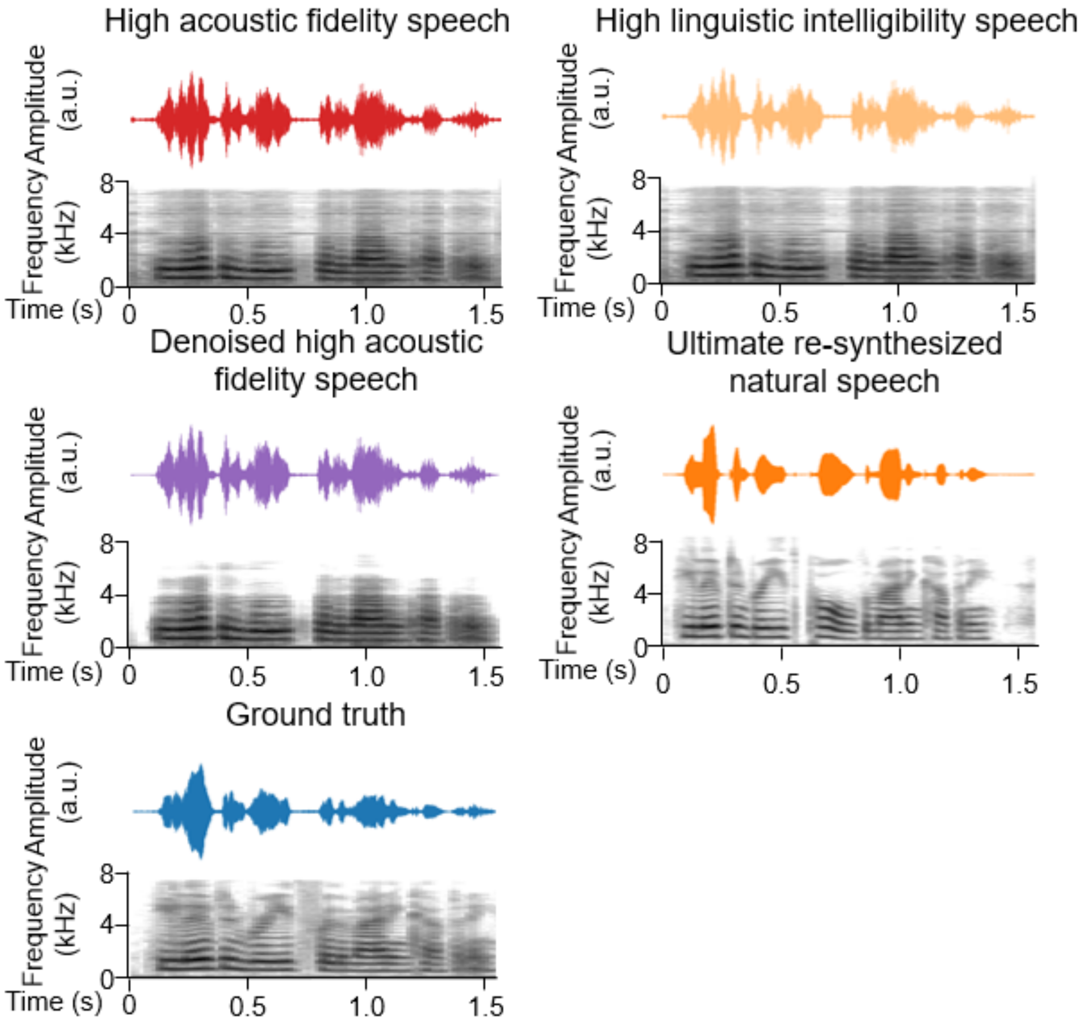


**Supplementary Figure 2. An example of original and re-synthesized speech in all stages.**

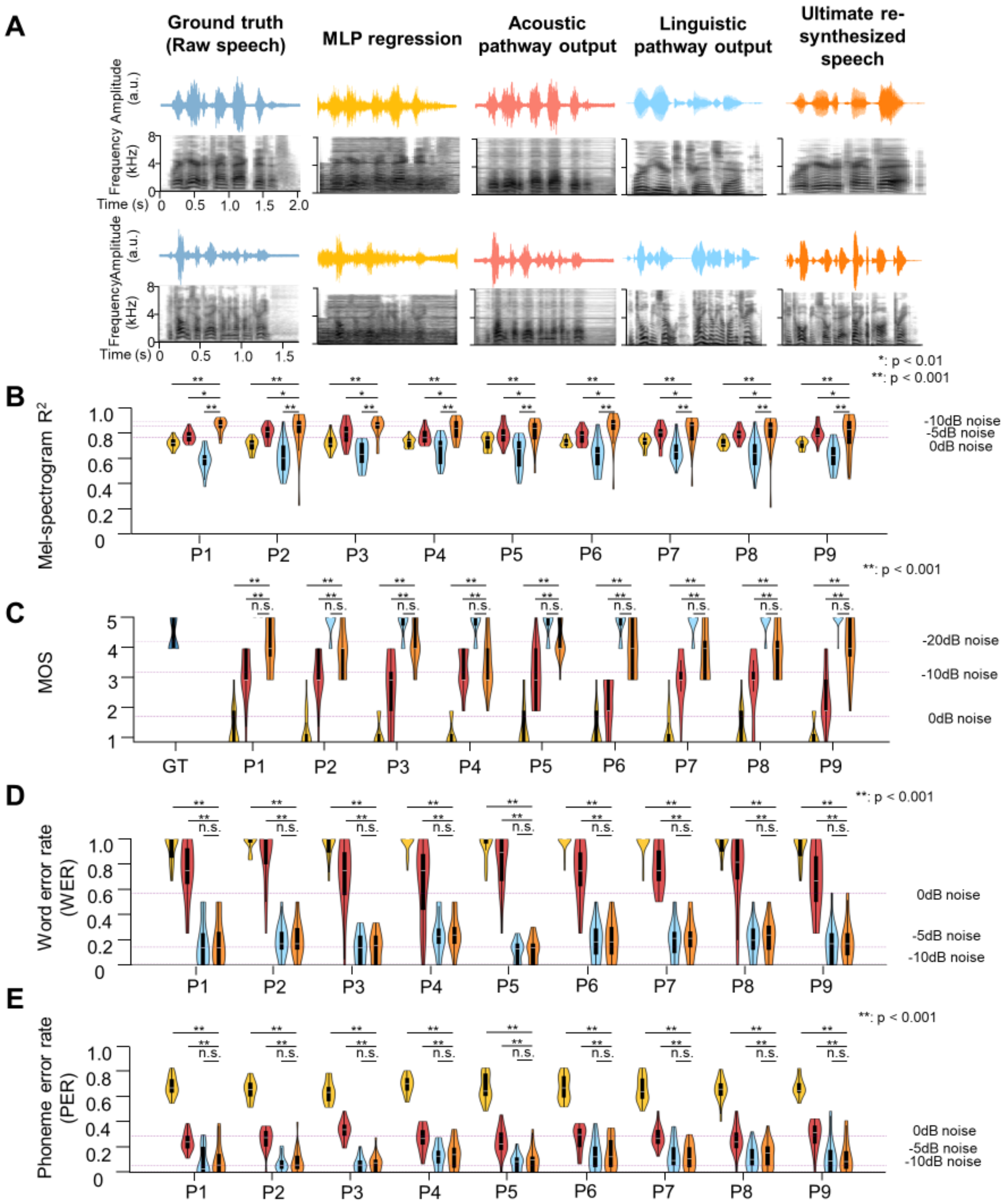


**Supplementary Figure 3 | Performance comparison between re-synthesized speech and baselines for each participant**

(**A**). Speech waveform and mel-spectrogram observations. Depicted are waveforms (time vs. amplitude) and mel-spectrograms (time vs. frequency ranging 0-8 kHz) for illustrative speech samples (Ground truth, baseline 1-3, and our ultimate-re-synthesized natural speech).

(**B**). Objective evaluation using mel-spectrogram $R^2$. Violin plots show the distribution of $R^2$ scores (0-1 scale) assessing spectral fidelity. The white dot represents the median, the box spans the interquartile range (25th to 75th percentiles), and whiskers extend to +/− 1.5 times the interquartile range, and the violin width illustrates data density at each point on the y-axis. The three shades of dashed lines from light to dark represent the average mel-spectrogram $R^2$ after adding additive noise at -10 dB, -5 dB, and 0 dB to the original speech waveform. The arrow on the right represents the direction of better results. P1-P9: participants.

(**C**). Subjective quality evaluation using mean opinion score (MOS). Violin plots show the distribution of MOS ratings (1-5 scale) assessing speech quality. The white dot represents the median, the box spans the interquartile range (25th to 75th percentiles), and whiskers extend to +/− 1.5 times the interquartile range, and the violin width illustrates data density at each point on the y-axis. The three shades of dashed lines from light to dark represent the average MOS scores after adding additive noise at -20 dB, -10 dB, and 0 dB to the original speech waveform. The arrow on the right represents the direction of better results. GT: Ground truth (original speech). P1-P9: participants.
(**D**). Intelligibility assessment using word error rate (WER). Violin plots show the distribution of WER scores (0-1 scale) assessing speech recognition accuracy. The white dot represents the median, the box spans the interquartile range (25th to 75th percentiles), and whiskers extend to +/− 1.5 times the interquartile range, and the violin width illustrates data density at each point on the y-axis. The three shades of purple lines from light to dark represent the average WER scores after adding additive noise at -10 dB, -5 dB, and 0 dB to the original speech waveform. The arrow on the left represents the direction of better results (lower WER). P1-P9: participants.
(**E**). Similar to panel (D), but evaluated using phoneme error rate (PER).

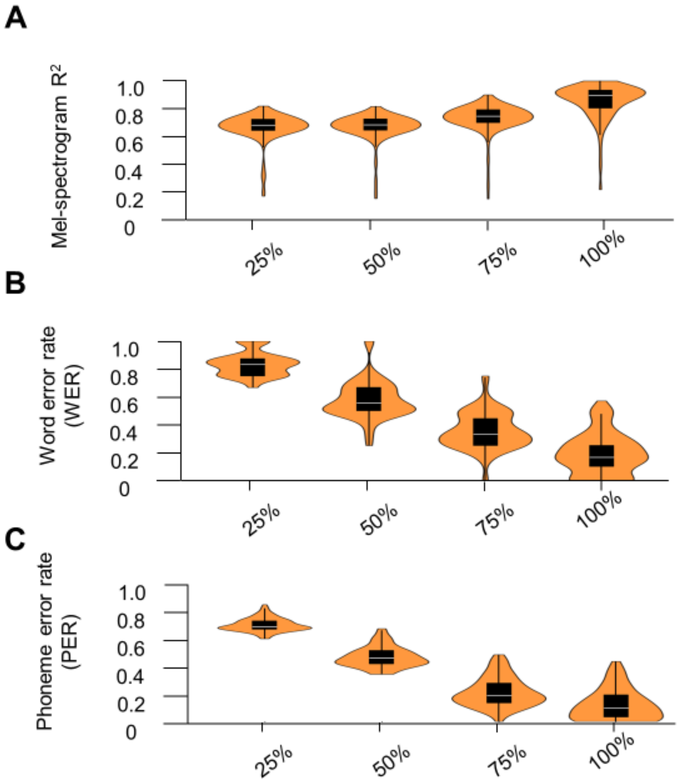


**Supplementary Figure 4 |**. Model performance scales with the amount of training data. (**A**) Mel-spectrogram correlation ($R^2$) between reconstructed and original speech. (**B**) Word Error Rate (WER). (**C**) Phoneme Error Rate (PER). All metrics are shown under the percentages of the used training set (25%, 50%, 75%, and 100%)

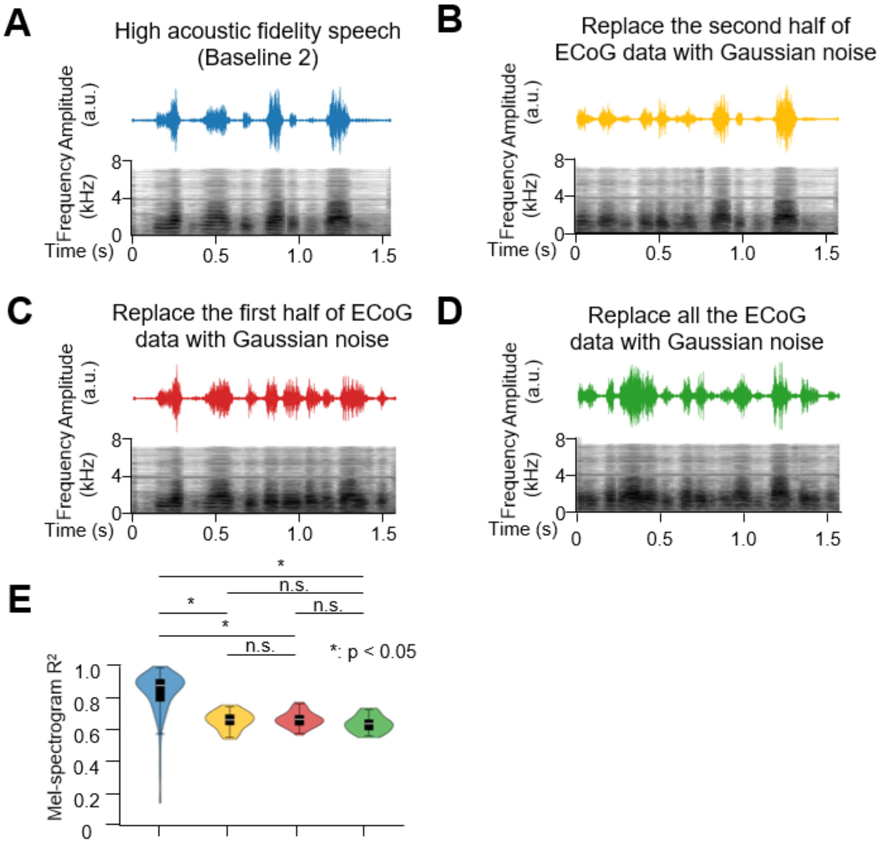


**Supplementary Figure 5. | Control analysis: Sensitivity of the acoustic pathway's reconstruction to neural input.**

(**A**) High-fidelity speech waveform (top) and corresponding mel-spectrogram (bottom) reconstructed from the full, veridical ECoG input by the acoustic pathway (Baseline 2).

(**B-D**) Reconstructions when portions of the input ECoG signal are replaced with Gaussian noise of matched dimensionality. (B) Replacing the first half of the ECoG signal with noise. (C) Replacing the second half of the ECoG signal with noise. (D) Replacing the entire ECoG signal with noise.

(**E**) Quantitative comparison of acoustic fidelity. Bar plot shows the mel-spectrogram $R^2$ values (mean ± s.e.m. across test samples) for the four conditions depicted in panels A-D: (1) Full ECoG input (A), (2) First half replaced with noise (B), (3) Second half replaced with noise (C), and (4) Entire signal replaced with noise (D). The significant drop in $R^2$ for conditions involving noise replacement confirms that reconstruction quality is causally dependent on and temporally locked to the veridical neural signal.

| Adaptor | Layers | Mel-spectrogram $R^2$ |
|---|---|---|
| LSTM | 1 | 0.780 ± 0.016 |
| LSTM | 2 | 0.788 ± 0.016 |
| LSTM | 3 | 0.793 ± 0.016 |
| LSTM | 4 | 0.789 ± 0.015 |
| LSTM | 5 | 0.792 ± 0.015 |
| Transformer | 1 | 0.759 ± 0.016 |
| Transformer | 2 | 0.761 ± 0.016 |
| Transformer | 3 | 0.761 ± 0.017 |
| Transformer | 4 | 0.757 ± 0.017 |
| Transformer | 5 | 0.727 ± 0.014 |

**Supplementary Table 1. Ablation study on adaptor architecture for the acoustic pathway.** Performance is evaluated by the mel-spectrogram $R^2$ (mean ± s.e.m. across participants), measuring the fidelity of reconstructed acoustic features. The bidirectional LSTM adaptor consistently outperformed the Transformer-based adaptor across different layer depths. The optimal performance was achieved with a 3-layer LSTM, which was selected for the final model.

| Adaptor | Layers | WER (%) | PER (%) |
|---|---|---|---|
| LSTM | 1 | 28.0 ± 4.0 | 15.9 ± 2.7 |
| LSTM | 2 | 26.7 ± 4.0 | 15.0 ± 2.7 |
| LSTM | 3 | 25.0 ± 3.2 | 14.1 ± 2.5 |
| LSTM | 4 | 24.8 ± 3.6 | 14.0 ± 2.5 |
| LSTM | 5 | 25.7 ± 4.0 | 14.4 ± 2.6 |
| Transformer | 1 | 22.5 ± 3.8 | 12.7 ± 2.4 |
| Transformer | 2 | 19.9 ± 3.4 | 11.6 ± 2.3 |
| Transformer | 3 | 17.7 ± 3.2 | 11.0 ± 2.3 |
| Transformer | 4 | 19.9 ± 3.4 | 11.5 ± 2.3 |
| Transformer | 5 | 21.3 ± 3.6 | 12.2 ± 2.3 |

**Supplementary Table 2. Ablation study on adaptor architecture for the linguistic pathway.** Performance is evaluated by word error rate (WER) and phoneme error rate (PER) (mean ± s.e.m. across participants), measuring the intelligibility of reconstructed speech. The Transformer-based adaptor achieved lower error rates than the LSTM-based adaptor across nearly all layer configurations. The optimal performance was achieved with a 3-layer Transformer, which was selected for the final model.

| Authors | Year | Neural recording modality | Neural recording durations | Primary task | Decoding framework | Performance |
|---|---|---|---|---|---|---|
| Akbari et al. *(1)* | 2019 | ECoG | 30 minutes | Speech perception | CNN + Vocoder | MOS=3.4<br>0.35 < ESTOI < 0.40 |
| Komeiji et al. *(2)* | 2022 | ECoG | 10-15 minutes | Speech production | CNN + Transformer + LSTM | PER = 31.3% |
| Bellier et al. *(3)* | 2023 | ECoG | 190.72 seconds | Music perception | MLP | Spectrogram $R^2$ = 0.429 |
| Willett et al. *(4)* | 2023 | Utah array | Approximately 100 hours | Speech production | GRU | WER = 24.7%<br>PER = 20.9% |
| Metzger et al. *(5)* | 2023 | ECoG | 20.8 hours | Speech production | RNN + pre-trained speech encoder | WER = 25.5% |
| Li et al. *(6)* | 2024 | ECoG | 20 minutes | Speech perception | LSTM + pre-trained speech decoder | ESTOI = 0.371<br>PER = 28.6%<br>MOS = 2.9 |
| Chen et al. *(7)* | 2024 | ECoG | 200 seconds | Speech perception and production | ECoG decoder + pre-trained speech synthesizer | Spectrogram $R^2$ = 0.81 |
| Wairagkar et al. *(8)* | 2025 | Utah array | More than 400 days | Online speech production | Transformer + TTS model | Spectrogram $R^2$ = 0.83<br>WER = 45.8%<br>PER = 34.0% |
| Li et al. (Ours) | 2025 | ECoG | 20 minutes | Speech perception | Acoustic and linguistic pathways (adaptor + generator) + voice cloner | Mel-spectrogram $R^2$ = 0.82<br>MOS = 4.0<br>WER = 18.9%<br>PER = 12.0% |

**Supplementary Table 3. Comparative overview of recent studies in neural-driven speech synthesis and re-synthe.**

The table summarizes representative work, highlighting the neural recording modality, approximate amount of data used for decoder training per subject, the primary experimental task (perception or production), and reported performance metrics. Studies are ordered chronologically. Performance metrics include: Mean Opinion Score (MOS, scale 1-5), Extended Short-Time Objective Intelligibility (ESTOI, scale 0-1), Word Error Rate (WER, %), Phoneme Error Rate (PER, %), and mel-spectrogram correlation ($R^2$, scale 0-1). Note that direct numerical comparisons should be made with caution due to differences in neural signals, tasks, stimuli, and evaluation methodologies across studies. Our study (highlighted in bold) achieves a competitive balance between data efficiency (~20 minutes) and performance across multiple metrics (WER, PER, MOS, $R^2$).
